# Toxicological profile of Umbilical Cord Blood-derived small Extracellular Vesicles

**DOI:** 10.1101/2021.06.30.450343

**Authors:** S.C. Rodrigues, R.M.S. Cardoso, C.F. Gomes, F.V. Duarte, P. C. Freire, R. Neves, J. Simões-Correia

## Abstract

The development and adoption of cell therapies has been largely limited by difficulties associated with their safety, handling and storage. Extracellular vesicles (EV) have recently emerged as a likely mediator for the therapeutic effect of cells, offering several advantages over cell therapies. Due to their small size and inability to expand and metastasize, EV are generally considered safer than cell transplantation. Nevertheless, few studies have scrutinized the toxicity profile of EV, particularly after repeated high dose administration. The present study aimed to evaluate a preparation of small EV obtained from umbilical cord blood mononuclear cells (UCB-MNC-sEV) for its cytotoxicity in different cell lines, as well as its differential accumulation, distribution and toxicity following repeated intravenous (IV) administrations in a rodent model. *In vitro*, repeated sEV exposure in concentrations up to 1×10^11^ particles/ml had no deleterious impact on the viability or metabolic activity of peripheral blood mononuclear cells, THP-1 monocytes, THP-1-derived macrophages, normal dermal human fibroblasts or human umbilical vein endothelial cells. DiR-labeled sEV, injected IV for four weeks in healthy rats, were detected in clearance organs, particularly kidneys, spleen and liver, similarly to control dye. Moreover, repeated administrations during six and twelve weeks of up to 1×10^10^ total particles of sEV-dye were well tolerated, with no changes in general hematological cell counts, or kidney and liver toxicity markers. Importantly, unlabeled sEV likewise did not induce significant alterations in cellular and biochemical blood parameters, nor any morphological changes in heart, kidney, lung, spleen, or liver tissue. In sum, our data shows that UCB-MNC-sEV have no significant toxicity *in vitro* or *in vivo*, even when administered repeatedly at high concentrations, therefore confirming their safety profile and potential suitability for future clinical use.

## Introduction

Extracellular vesicles (EV), secreted by most cell types, are described as key mediators of intercellular communication, through the transport of a wide array of bioactive molecules, such as proteins, RNA (including microRNAs), and DNA^1,2^. Being such a heterogeneous group, EV are catalogued in different subsets according to their size, ranging from the micron to the sub-micron dimension^3,4^. This diverse group of biological carriers participates in various physiological and pathophysiological processes, which sparked the scientific interest in recent years. Due to their biological and functional roles, EV have emerged as potential candidates for the replacement of cell therapies in different diseases contexts^5,6^. In regenerative medicine, EV isolated from mesenchymal stromal cells (MSC) and mononuclear cells (MNC) have been demonstrated to successfully replace cell-based therapies, improving the function of damaged organs in animal models of ischemic diseases, such as stroke^7,8^, cardiovascular diseases^6,9,10^ and chronic wounds^11,12^.

As a promising new tool in regenerative medicine, several considerations must be taken before its clinical use. A major attribute of a new medicine before its commercialization relies in the establishment of its biological safety through appropriate toxicological studies^13^. In fact, intravascular infusion of MSC has been documented to cause embolism and death in a mouse model^14^, whereas MSC inoculated into pig infarcted myocardium were reported to induce adverse cellular growth such as cardiac sympathetic nerve sprouting^15^. For adverse effects such as these, it appears likely that the risk associated with EV administration, due to their small sizes, will be significantly lower or perhaps absent^16^. Moreover, EV are less prone to trigger immune responses and are unable to directly form tumors. A recent study by Zhu et al. reported minimal toxicity and immunogenicity of HEK293T-derived EV following repeated dosing in C57BL/6 mice^17^. Similarly, EV isolated from suspension human embryonic kidney Expi293F cells showed minimal toxicity and pro-inflammatory cytokine response following systemic administration into BALB/c mice^18^. However, the diversity of sources, isolation protocols and manipulation of EV makes it difficult to transversally accept this claim for all EV-based therapies. Thus, it is increasingly urgent and fundamental to recommend standard techniques for the clinical grade production and quality control of EV-based therapies, as well as the definition of toxicology studies to accurately assess the safety of these new therapies.

In this study, EV isolated from human umbilical cord blood mononuclear cells (UCB-MNC) through a clinically transferable process were tested for their toxicity *in vitro* and *in vivo*. As described in a recent paper, the optimized methodology combines ultrafiltration and size exclusion chromatography (UF/SEC), yielding a small EV (sEV)-enriched product, with particle sizes ranging from 50-200 nm^19^. In comparison with ultracentrifugation, UF/SEC significantly reduces production time and improves process standardization^20,21^, while maintaining sEV’s bioactivity^19^. This optimization in the manufacturing process may increase the confidence in UCB-MNC-sEV’s safety due to their strictly controlled production process. However, we cannot predict if these sEV induce metabolic or cell viability alterations, bring inflammatory and immune responses or hematologic variations.

UCB-MNC-sEV preparations consist of 80% sEV and 20% larger vesicles, as well as proteins and lipids^19^. So far, intravenous injection of similar EV showed no signs of toxicity^18^. The present study aimed to evaluate the cytotoxicity of UCB-MNC-sEV *in vitro*, as well as determine their differential accumulation, distribution and toxicity following repeated intravenous injection in a rodent model.

## Material and methods

### UCB-MNC-sEV isolation and purification

Human UCB samples were obtained upon signed informed consent, in compliance with Portuguese legislation. The collection was approved by the ethical committee of Centro Hospitalar e Universitário de Coimbra, Portugal. Samples were stored and transported to the laboratory in sterile bags with anticoagulant solution (citrate-phosphate-dextrose) and processed within 48 hours after collection. UCB units were processed in an accredited cryobank (Crioestaminal, Cantanhede, Portugal), using an automated system AXP, according to the manufacturer’s recommendations.

After at least one week of storage, UCB-MNC were thawed and cultured at 2 million cells/mL in X-VIVO 15 serum-free cell-culture medium (Lonza, Basel, Switzerland) supplemented with 0.5 μg/mL of FMS-like tyrosine kinase-3 and 0.5 μg/mL of stem-cell factor, under ischemia (0.5% O2) conditions. Following 18 hours of secretion, conditioned media were cleared by centrifugation and filtration, followed by ultrafiltration at 3-bar with a 100 kDa filter (Sartorius, Goettingen, Germany). Finally, the concentrated conditioned medium underwent size exclusion chromatography and EV-enriched fractions were collected, concentrated, and stored at -80°C until further use. A more detailed description of the EV purification process is published elsewhere (19).

### UCB-MNC-sEV morphological characterization

Size distribution and concentration of UCB-MNC-sEV was measured in Nanosight LM (Malvern Instruments Ltd, Malvern UK) equipped with 638 nm laser and a CCD camera. The measurements were performed with a detection threshold set to 3, camera level set to 13 and a screen gain of 10. The blur and Max Jump Distance were set to 2. The sEV samples were diluted to obtain a number of particles per frame between 15 and 30. Readings were taken in 5 captures during 30 sec each at manual monitoring of temperature.

### In vitro toxicity

To evaluate the possible cytotoxic effects of UCB-MNC-sEV *in vitro*, three UCB-MNC-sEV concentrations (1×10^10^, 5×10^10^ and 1×10^11^ particles/ml) were tested on different cell types. After 72 hours of incubation, cell viability was evaluated through an XXT assay. PBS was used as vehicle control and all experiments were performed using EV-depleted medium.

#### Cell Culture

Normal human dermal fibroblasts (NHDF) and human umbilical vein endothelial cells (HUVEC) (ATCC, VA, USA) were cultured in T75 culture flasks, maintained in an incubator with controlled temperature (37 °C), 5% of CO2 level and 90% humidity. 60,000 cells were plated in 96-well plate to the XTT assay. After 24h of plating the cells and immediately before treatment, the culture medium was replaced by EV-depleted medium (previously centrifuged for 14h at 100,000xg).

THP-1 cells (monocytes; ATCC) were cultured in T75 culture flasks, maintained in an incubator with controlled temperature (37 °C), CO2 level (5%). These cells were grown to a density of 7 x 10^5^ cells/ml, plated in a 96-well plate in EV-depleted medium and rested in culture 24 hours before treatment. To obtain macrophages, 100,000 THP-1 monocytes were seeded and incubated for 48 hours with 25 nM PMA. Then, the PMA-medium was replaced by fresh medium and adherent cells were rested in culture 24 additional hours. Immediately before treatment, the culture medium was replaced by EV-depleted medium.

Human blood samples were obtained from Hospital Universitário de Coimbra, where donations were obtained from healthy volunteers after providing their informed consent. Peripheral blood mononuclear cells (PBMC) were isolated by Lymphoprep gradient centrifugation (Stemcell Technologies, Vancouver, Canada) and plated on 96-well flat bottom culture plates (Corning-Costar, Milan, Italy) at a density of 10^5^ cells/well.

#### Cell viability - XTT Assay

To determine the viability of cells treated with UCB-MNC-sEV, an XTT assay (Applichem) was performed according to the supplier’s instructions. Briefly, XTT mix was added to the medium and incubated at 37 °C for 3.5 hours. The absorbance was read at 450nm and 630nm.

### Animal experiments

Animal testing protocols with Wistar Rats were approved by the Portuguese National Authority for Animal Health (DGAV). All surgical and necropsy procedures were performed according to the applicable national regulations, respecting international animal welfare rules.

#### In vivo Biodistribuition and toxicity: 4 weeks

Male Wistar Rats (12-week-old), purchased from Charles River and weighing between 250-400g, were housed in a specific pathogen-free animal facility on a 12 hours light/12 hours dark regimen and fed a commercial diet (pellets) and acidified drinking water *ad libitum*.

To assess the biodistribution of systemically delivered UCB-MNC-sEV in rats, the vesicles were labelled with DiR (C18) dye (Thermo Fisher Scientific, MA, USA). Briefly, 50 μM of DiR dye were incubated with UCB-MNC-sEV or added to vehicle (PBS) for 30 min at 37 °C. Then, the samples were ultracentrifuged at 100,000 xg for 2h18min at 4°C and filtered (0.2µM). Before use, labelled-sEV were analyzed by nanoparticle tracking analysis (NTA). After UCB-MNC-sEV modification with DiR dye, 50μl (5×10^10^ particles/mL) and the respective control were injected intravenously in rats tail vein twice a week. After four weeks of treatment, the fluorescent signal was observed with IVIS Lumina XR equipment (Caliper Life Sciences, Hopkinton MA).

During the experiment, the animals’ weight and wellbeing was monitored. Signals of fighting, dehydration, excessive barbering or malocclusion were closely monitored. After the 4 weeks, the animals were euthanized through recommended anesthetic overdose of xylazine and ketamine. The most relevant organs, namely liver, lungs, kidneys, spleen, pancreas and heart, were collected for further analysis of the fluorescence signal. Blood was also collected for further hemogram, leucogram and biochemistry analysis. Urea, creatinine, alanine aminotransferase (ALT), aspartate aminotransferase (AST) and alkaline phosphatase (ALP) biochemical analyses were performed in *Beatriz Godinho-Análises Clinicas* accredited laboratory. Several important hematology markers including leukocytes, neutrophils, eosinophils, basophils, lymphocytes, monocytes, red blood cell count (RBC), hemoglobin, hematocrit, mean corpuscular volume (MCV), cell haemoglobin (CH), mean cell haemoglobin concentration (CHCM), and RBC distribution width (RDW) were selected for further toxicity assessment of UCB-MNC-sEV *in vivo*.

#### In vivo Toxicity: 6 and 12 weeks

Male Wistar Han Rats (12-week-old), purchased from Charles River and weighing between 250-400g, were housed in a specific pathogen-free animal facility on a 12h light/12h dark regimen and fed a commercial diet (pellets) and acidified drinking water *ad libitum*.

A total of 40 animals were randomized into 3 groups: Group 1-UCB-MNC-sEV 1×10^10^ particles/mL (1×10^9^ total particles) (14 animals) ; group 2-UCB-MNC-sEV 1×10^11^ particles/mL (1×10^10^ total particles) (14 animals); group 3-vehicle (12 animals). Each group was randomly sub-divided into 2 groups depending on the duration of treatment: 6 or 12 weeks. UCB-MNC-sEV (100μl at 1×10^10^ or 1×10^11^ particles/mL) or control (vehicle) were intravenously injected into the tail vein, twice a week, over 6 or 12 weeks. During the experiment, the animals’ weight was monitored, and signals of fighting, dehydration, excessive barbering or malocclusion were closely observed. After 6 or 12 weeks, animals were euthanized and organs and blood were collected as described above.

#### Histological Examination

Tissue biopsies were formalin-fixed in neutral buffered formalin, paraffin-embedded, cut in 4 μm sections, and stained with hematoxylin (Bio-Optica, Milan, Italy) and eosin (Thermo Fisher Scientific, MA, USA). Tissue sections were analyzed by a pathologist blinded to experimental groups, in a Leica DM2000 microscope coupled to a Leica MC170 HD microscope camera (Leica Microsystems, Wetzlar, Germany).

### Statistical Analysis

Data were analyzed using GraphPad Prism 6 software. Statistical analysis was performed by unpaired t-test. The statistical significant level chosen was p-value (p)<0.05. Results were shown as mean ± standard error of the mean (SEM) and, when appropriated, they are marked with one asterisk (*) if p<0.05, ** for p<0.005, *** for p<0.0005, **** for p<0.0001 and non-significant (ns) if p>0.05.

## RESULTS

### In vitro toxicity

Intravenous route of administration not only allows avoidance of variability among absorption sites, but also represents a potential exposure route of UCB-MNC-sEV for their therapeutic applications. Upon intravenous injection, the first cells to contact with the testing material are blood cells. Then, upon distribution in the organism by systemic circulation, the compound reaches different organs passing through capillaries and connective tissue. Hence, we aimed to evaluate cytotoxicity using primary cells and cell lines, challenging them with different sEV particle concentrations. Three UCB-MNC-sEV concentrations (1×10^10^, 5×10^10^ and 1×10^11^ particles/ml) were tested on blood/immune system cells (monocytes and PBMCs), endothelial cells (HUVECs) and fibroblasts (NHDF). These concentrations were chosen based on previous efficacy tests which demonstrated 1×10^10^ particles/ml as the optimal concentration to use. Therefore, 5- and 10-times higher concentrations were also included in this work. UCB-MNC-sEV were applied twice a day for 3 days. 72h after treatment, cell viability was evaluated by XTT assay. No evidence of cytotoxicity was found in any of the tested concentrations **(Figure 1)**. Namely, the metabolic activity of total PBMC, THP-1 monocytes and macrophages, NHDF and HUVEC was similar or increased, compared with control. Of note, the metabolic activity of total PBMC remained constant when stimulated with sEV concentrations up to 1×10^11^ particles/mL. Since an increased metabolic activity of these cells could indicate immunogenicity by UCB-MNC-sEV, we conclude that, under these *in vitro* conditions, UCB-MNCs-sEVs do not induce an immunogenic response on PBMCs.

**Figure 1.**
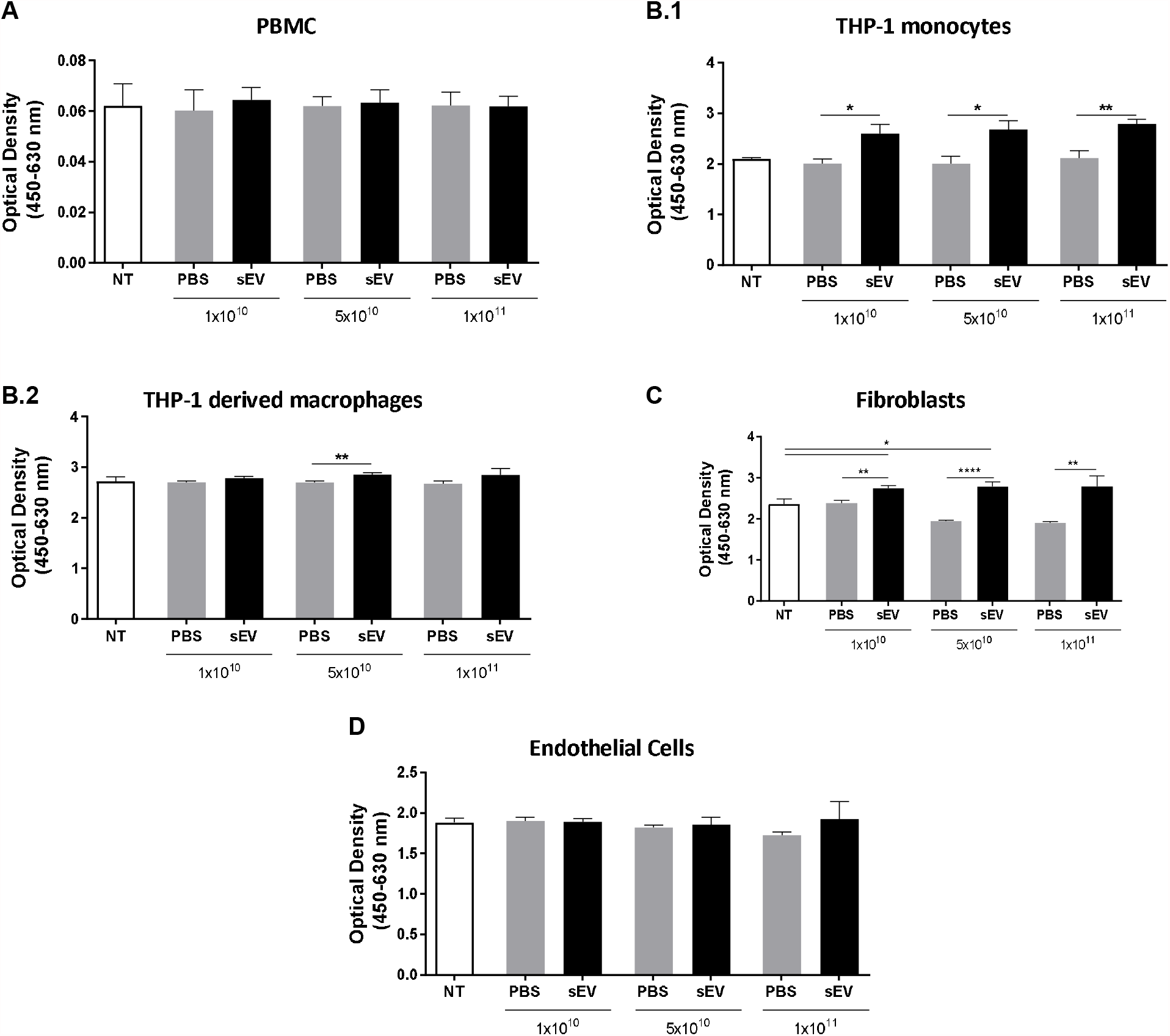
UCB-MNC-sEV do not induce cytotoxic effects *in vitro*. After cell seeding, cells received 6 doses of UCB-MNC-sEV with the indicated particle concentration. After a 72h treatment, cellular metabolic activity was was measured by XTT assay on (A) PBMC (n≥3 per condition), (B.1) THP-1 monocytes (n≥3 per condition), (B.2) THP-1 derived macrophages (n≥3 per condition), (C) fibroblasts (n≥6 per condition) and (D) endothelial cells (n≥6 per condition). NT refers to “not treated” conditions. Results are presented as mean ± SEM. Statistical analysis was performed by unpaired t-test, * for p-value<0.05, ** for p-value<0.005 and **** for p-value<0.0001.

In contrast, UCB-MNC-sEV administration on a monocytic cell line (THP-1) increased cellular metabolic activity. This increase is similar across the three UCB-MNC-sEV tested concentrations (≈31 %). As XXT activity is also a proliferation readout, this increase could be due to a high proliferation index on sEV-treated conditions. Interestingly, when the same cell line was differentiated into macrophage-like cells by PMA induction, a different pattern was observed. In this case, only 5×10^10^ particles/mL concentration of UCB-MNC-sEV induced a marginal increase on cell metabolic activity (≈6%).

Similarly, the results obtained in fibroblasts demonstrated a significant increase in cell metabolic activity, as measured with XTT assay **(Figure 1C**) in all tested concentrations. The greatest difference was observed upon treatment with 5×10^10^ concentration of UCB-MNCs-sEVs (≈43%). Accordingly to what was previously reported by Antunes et al., UCB-MNC-sEV are able to promote fibroblast migration and proliferation^12^.

In an endothelial cell line, no significant differences were detected between control and tests groups.

In sum, UCB-MNC-sEV do not elicit any cytotoxic effect *in vitro* when used between 1×10^10^ and 1×10^11^ particles/mL.

### In vivo biodistribuition and toxicity: 4 weeks

To confirm previous *in vitro* findings in an *in vivo* model, we first assessed the biodistribution and bioaccumulation of the UCB-MNC-sEV. Wistar Han Rats were used and treated with dye-modified UCB-MNC-sEV in solution (5×10^10^ particles/mL) by tail vein injection, twice a week, over 4 weeks. The main organs were collected and analyzed by IVIS.

Data extracted from fluorescence accumulation showed that after 4 weeks of treatment, dye-modified UCB-MNC-sEV and control were mostly accumulated in kidneys, followed by the spleen and liver **(Figure 2A and B)**. However, the accumulation effects cannot be attributed exclusively to sEV, as the respective control (dye without EV) is distributed equally among rat organs, with no significant differences noted between the two groups.

**Figure 2.**
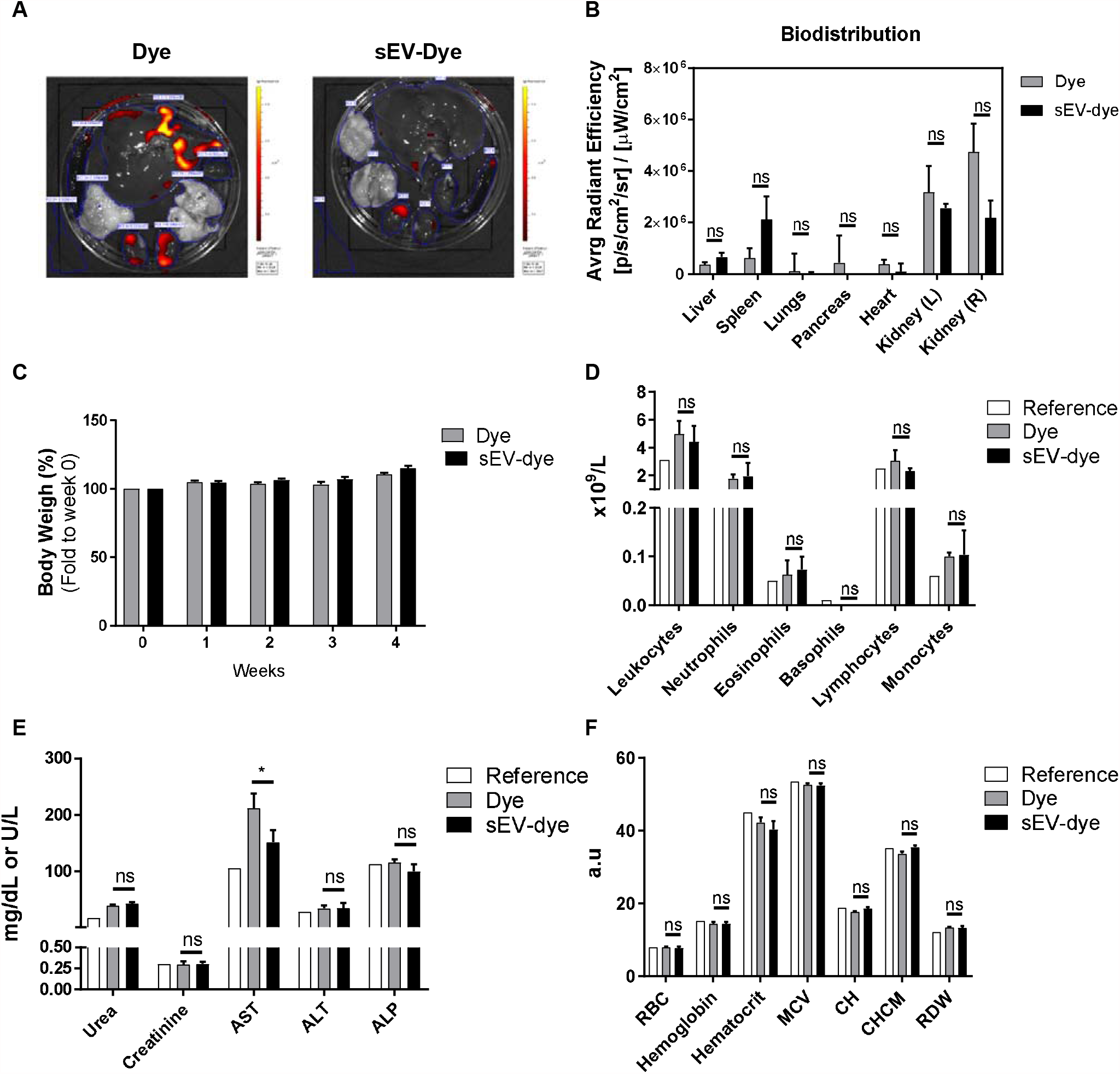
Biodistribution and toxicological profile of UCB-MNCs-sEVs in rats. Wistar Han rats were injected intravenously with fluorescently labelled UCB-MNCs-sEVs in PBS, twice a week for 4 weeks. After sacrifice, organs were analyzed for accumulated fluorescence (IVIS) and blood was collected to evaluate signs of systemic toxicity. Control (Dye) animals were injected with the same dye concentration as used for labeled UCB-MNCs-sEVs. (A-B). (C) Graphic representation of weight evolution during the 4-week experiment. Graphic representation of (D) leukogram, (E) biochemical and (F) hemogram results acquired at the end of the experiment. Reference values were obtained from Charles River. Results are presented as mean ± SEM. Statistical analysis was performed by using unpaired t-test. (n=4 rats per condition). RBC: red blood cell count; MCV: mean corpuscular volume (MCV); CH: cell haemoglobin; CHCM: mean cell haemoglobin concentration; RDW: RBC distribution width; a.u.: arbitrary units. ns, non-significant; *p-value<0.05.

As shown in **Figure 2C-D**, there was no significant impact on rats’ weight or circulating cell populations during the experiment, for either dye-sEV or dye alone. Aspartate aminotransferase (AST), an enzyme indicative of liver damage when found in the circulation, was significantly higher in rats receiving dye alone versus animals dosed with dye-sEV (Figure 2E). Given that this result was not accompanied by other markers of liver damage, it likely does not represent any significant tissue damage. Still, animals receiving dye-sEV showed normal levels of AST.

Finally, a classic hemogram analysis showed no relevant differences between reference values and the two test groups (Figura 2F).

### In Vivo Toxicity: 6 and 12 weeks

We next aimed to verify the bioaccumulation effects and toxicity of the UCB-MNC-sEV in a worst-case scenario study (high dose, intravenous route, and repeated-dose long-term application). For that, we used Wistar Han Rats, applying UCB-MNC-sEV in a saline solution (1×10^10^ and 1×10^11^ particles/mL, respectively 1×10^9^ and 1×10^10^ total particles) by tail vein injection, twice a week, over 6 or 12 weeks. The blood and main organs were collected and analysed. Results show that UCB-MNC-sEV have no significant impact on rats’ weight over time (**Figure 3A**). All animals showed an increase in body weight, suggestive of good general health and appropriate access to food and water.

**Figure 3.**
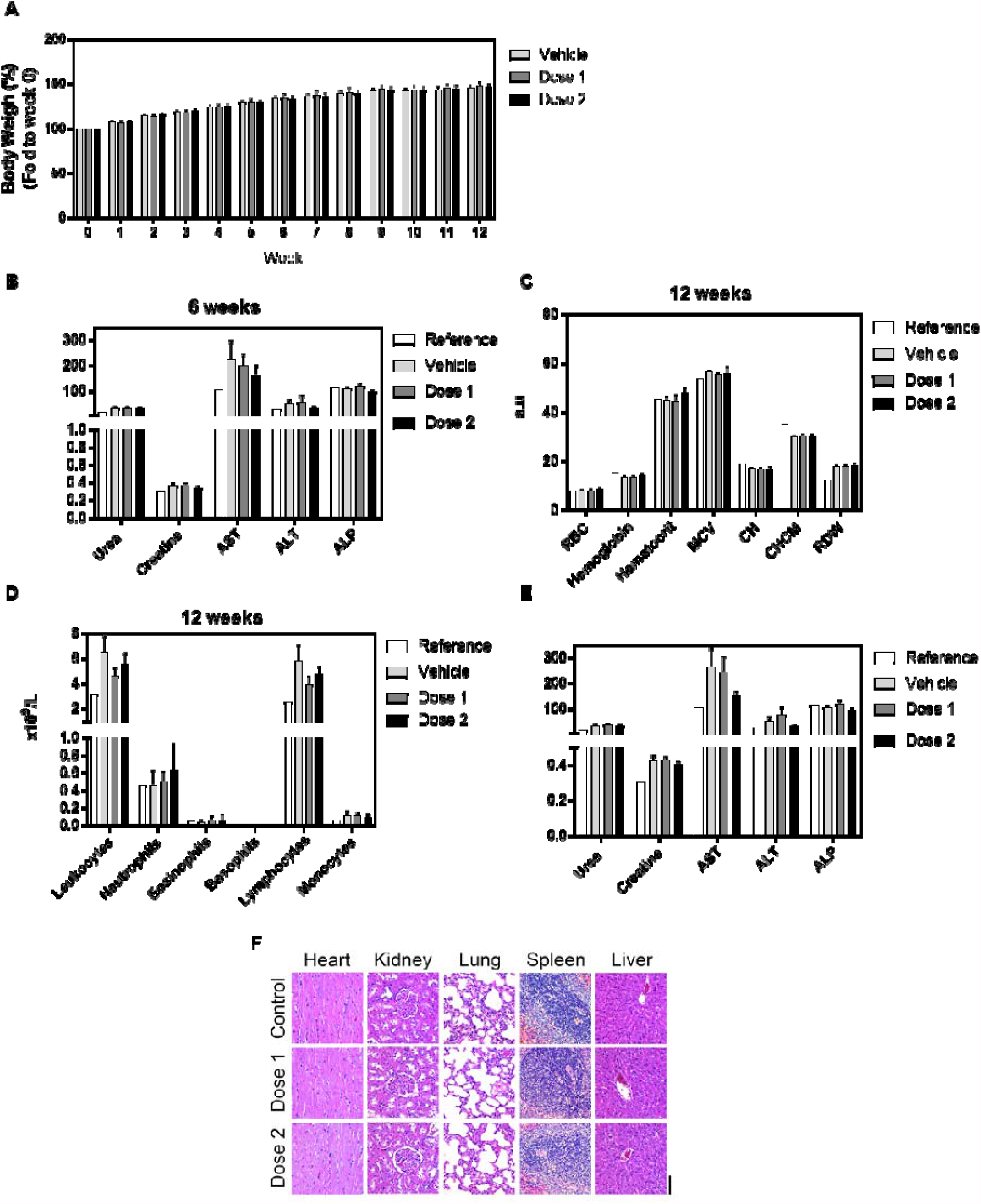
Toxicological profile of UCB-MNCs-sEVs in rats after repeated dose treatment (6 and 12 weeks). Wistar Han rats were injected intravenously with two UCB-MNCs-sEVs doses, twice a week for 6 or 12 weeks. Dose 1 and 2 refer to 1×10^10^ and 1×10^11^ particles/ml (1×10^9^ and 1×10^10^ total particles), respectively. After sacrifice, organs and blood were collected for analysis. Control (vehicle) animals were injected only with saline solution (PBS). (A) Graphic representation of weight evolution during the 12 weeks experiment. Graphic representation of (B) biochemical, (C) hemogram, (D) leukogram and (E) biochemical results acquired at the end of the experiment. Reference values were obtained from Charles River. Results are presented as mean ± SEM. Statistical analysis was performed by using unpaired t-test. (n=7 rats per condition). (F) Representative H&E microphotographs of heart, kidney, lung, spleen and liver from untreated and treated rats exposed to 2 different doses. Original magnification 20x (bar, 100µm).

Regarding the analyses performed on the animals’ blood, both in the 6- and 12-week group, it can be observed that there are no significant differences in any of the parameters evaluated (hemogram, leukogram and biochemical analyses), as shown in **Figure 3B-E**. We have conducted histological analysis of major functional organs, including heart, kidney, lung, spleen and liver, and observed no morphological changes or signs of toxicity in any of the organs analyzed (**Figure 3F**).

## Discussion and Conclusion

In this study, the toxicology of UCB-MNC-sEV was analysed to assess the vesicles’ feasibility as therapeutic agents and predict potential adverse effects. EV contain most of the desirable features of an ideal drug delivery system, such as the intrinsic ability to target tissues, biocompatibility, and presumably minimal toxicity ^22^. Nevertheless, it is imperative to know the toxicological risks of sEV before their clinical use^13^.

Collectively the results obtained provide evidence for the absence of significant toxicity after treatment with UCB-MNC-sEV. *In vitro*, the results show no UCB-MNC-sEV cytotoxicity, since there is no significant decrease in relative cell number or cell metabolic activity, as measured with an XTT assay. Notably, dermal fibroblasts and monocytic cells presented a significant increase in cell viability, which could be a cellular effect induced by UCB-MNC-sEV. It is widely accepted that sEV have the capacity of recruiting monocytes to injury sites and promoting the secretion of different anti-inflammatory biomolecules^23–25^. We believe that the metabolic boost observed in THP-1 cells is correlated with UCB-MNC-sEV’s potential to modulate inflammatory responses. On the other hand, multiple reports demonstrate EV’s potential to induce the proliferation of dermal cells, namely fibroblasts^26–28^. These outcomes are in line with previous UCB-MNC-sEV effects observed in wound healing, both in *in vitro* and *in vivo* models^12,19^.

Based on our results, we conclude that UCB-MNC-sEV do not elicit any severe immune response or toxicological effect. Due to the naïve nature of UCB-MNC, no immunogenicity was expected, although EV from different sources are able to trigger an intense immune response^29,30^. Similar reports of low immunogenicity are demonstrated when EV from UCB-MSC are used^31,32^. Overall, our *in vitro* data demonstrate that UCB-MNC-sEV do not significantly decrease cell metabolic activity of different cell types, at least within the range of concentrations tested, until after 72h of contact with sEV.

Biodistribution and bioaccumulation studies, after 4 weeks of treatment, revealed no significant differences in weight, hemogram and leucogram between the evaluated groups. The distribution and accumulation of UCB-MNC-sEV systemically delivered to rats was measured using sEV labelled with DiR dye and posteriorly detected by IVIS, as reported also by other authors^33–35^. Dye-modified UCB-MNC-sEV seem to accumulate more in the liver and spleen. This pattern of EV accumulation is consistent with previous reports examining EV biodistribution in mice ^33,34^. Nevertheless, these reports demonstrated an inverse order of fluorescence accumulation, being higher in liver followed by spleen, lung and brain^36^. Therefore, we believe in a preferential accumulation in spleen compatible with sEV’s immunomodulatory mode of action. Hepatic and splenic bioaccumulation is related to the uptake by resident phagocytes in the liver, as well as macrophages and B cells in the spleen, that are part of the mononuclear phagocytic system (MPS), or of the reticuloendothelial system (RES), as described in other biodistribution reports^33,37^. In addition, much of the signal detected by IVIS in both treated and control groups were noticed in kidneys. In view of the substantial variability and the absence of statistically significant data, consideration should be given to the hypothesis that the data obtained may have been influenced by the dye excretion pathway, and the obtained results do not represent the bioaccumulation pattern of UCB-MNC-sEV.

Considering the biochemistry analyses, the DiR dye seems to induce a slight hepatoxicity by increasing AST levels. Interestingly, when this dye is incorporated in sEV, AST levels were significantly decreased, in comparison with dye alone. These results suggest either a sEV hepatic protective mechanism or dye’s inability to interact with membranes once it is already conjugated with sEV. In addition, the histological analysis corroborates the absence of toxicity induced by UCB-MNC-sEV administration, since all sections analyzed depict normal histological features. Similarly, administration of Expi293F-derived EV to mice did not result in any histopathological changes or increases of liver transaminases, supporting minimal or absent liver damage ^18^.

To conclude, no harmful effects were caused by the intravenous injection of the highest tested dose of UCB-MNC-sEV. This dose is 40 times higher than the predicted therapeutic dose^12,19^. As this experiment was designed to resemble a worst-case scenario toxicity study, we can conclude that UCB-MNC-sEV use is safe at concentrations up to 1×10^11^ particles/mL (or 1×10^10^ total particles), when administered intravenously twice a week for three months. By confirming their safety in a rodent model, these results support the clinical development of UCB-MNC-sEV.

## Author contribution

R.M.S.C, S.C.R., C.F.G., F.V.D and J.S-C. conceived and designed experiments. R.M.S.C, S.C.R, C.F.G. and F.V.D performed experiments. R.M.S.C., S.C.R., C.F.G. and F.V.D. collected and analyzed data. R.M.S.C, S.C.R. and C.F.G. executed animal experiences. R.N. and J.S-C analyzed and reviewed data and the manuscript. C.F.G, S.C.R and P.C.F wrote the manuscript. All authors discussed the results, revised, and approved the manuscript.

## Acknowledgments

The work was co-funded by Regional Operational Program Center 2020, Portugal 2020 and European Union through FEDER within the scope of CENTRO-01-0247-FEDER-022398 and CENTRO-01-02B7-FEDER-070018, and supported by FCT fellowship SFRH/BD/137633/2018.

